# A benchmark of optimization solvers for genome-scale metabolic modeling

**DOI:** 10.1101/2023.04.11.536343

**Authors:** Daniel Machado

## Abstract

Genome-scale metabolic modeling is a powerful framework for predicting metabolic phenotypes of any organism with an annotated genome. For two decades, this framework has been used for rational design of microbial cell factories. In the last decade, the range of applications has exploded, and new frontiers have emerged, including the study of the gut microbiome and its health implications, and the role of microbial communities in global ecosystems. However, all the critical steps in this framework, from model construction to simulation, require the use of powerful linear optimization solvers, with the choice often relying on commercial solvers for their well-known computational efficiency. In this work, I benchmark a total of six solvers (two commercial and four open-source) and measure their performance to solve linear and mixed-integer linear problems of increasing complexity. Although commercial solvers are still the fastest, at least two open-source solvers show comparable performance. These results show that genome-scale metabolic modeling does not need to be hindered by commercial licensing schemes and can become a truly open science framework for solving urgent societal challenges.

## Introduction

Genome-scale metabolic modeling is a mathematical framework for predicting the metabolic phenotype of an organism based on genomic and environmental information^1^. Some of the first models were developed for industrially-relevant organisms like *Escherichia coli*^2^ and *Saccharomyces cerevisiae*^3^, and used to find optimal intervention strategies for rational strain design^4^. Recently, automated reconstruction tools^5,6^ have enabled the creation of large model collections, accounting for virtually any organism that has been sequenced. This has expanded the frontiers of research using genome-scale models to address most societal issues, from the study of the human gut microbiome and its health implications^7^, to global microbial ecosystems of the soils^8^ and oceans^9^.

While this research field exploded with new simulation methods and software tools10, one fundamental obstacle remained. This family of so-called constraint-based analysis and reconstruction (COBRA) methods requires efficient linear optimization problems that can handle thousands of variables and constraints. The choice has often fallen upon commercial optimization solvers like CPLEX (IBM) and GUROBI (Gurobi Optimization LLC) due to their known computational efficiency. The GNU Linear Programming Kit (GLPK) has been used as an open-source alternative, but it offers slower computation times and no longer seems to be actively maintained. This bottleneck in the selection of optimization solvers creates obstacles for users, who become restricted by complex licensing schemes, hindering genome-scale modeling from becoming a truly open science framework.

In this work, I benchmark the three solvers mentioned above together with three recent open-source solvers that can be a promising replacement for commercial solvers. The benchmark considers two main problem formulations: linear problems (LPs), which are commonly used to run flux balance analysis (FBA) simulations and other kinds of theoretical yield analysis^11^; and mixed-integer linear problems (MILPs), used to solve problems with any kind of selection constraints (implemented with binary or integer variables), such as optimal strain design^4^ or gap-filling^12^. Quadratic minimization problems were not considered as they are not supported by all solvers and can be reformulated as linear minimization problems (by replacing L2 with L1 norm).

## Methods

### Software setup

Six optimization solvers (Table 1) were installed in a local machine (8-core CPU with 16 GB of RAM). The open-source solvers were installed using the *conda* package manager. For the commercial solvers, software installers and free academic licenses were obtained from the respective websites. All solvers were executed with their default parameter configurations. The *PuLP* library^13^ (version 2.7.0) was used as a common interface to interact with all the solvers. All simulations were performed using the *ReFramed* library^18^ (version 1.3.0).

**Table 1:**
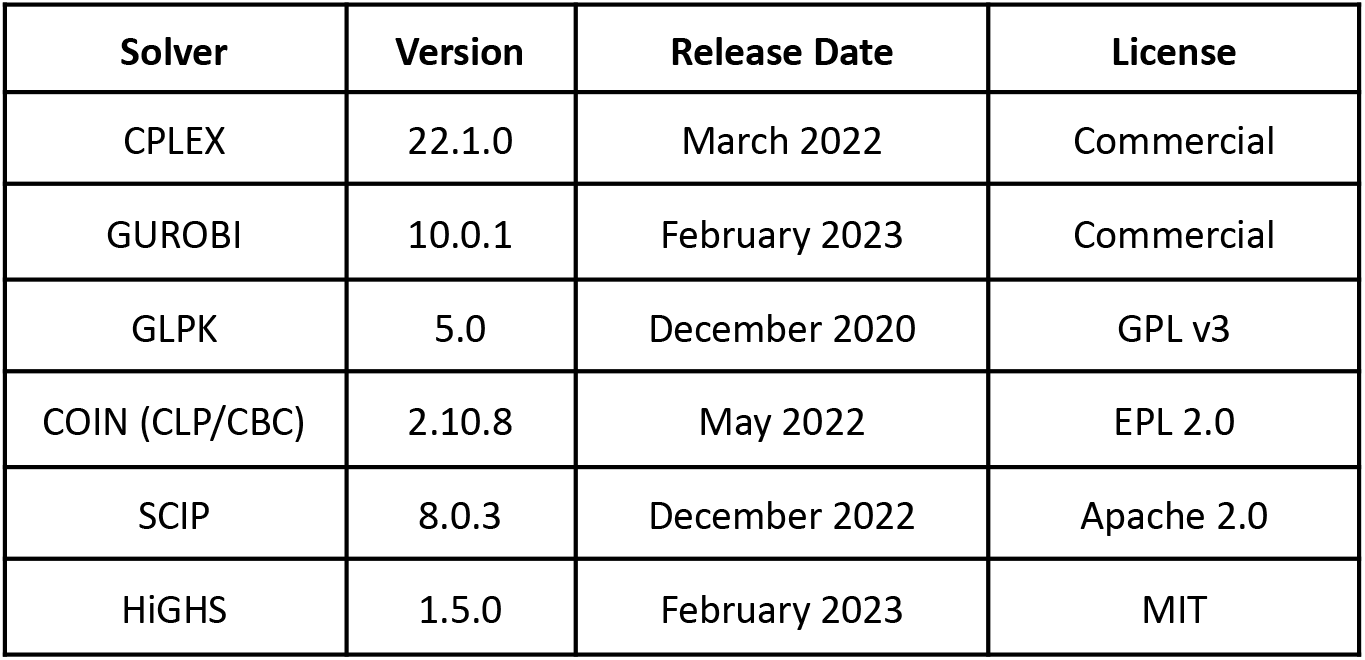
List of optimization solvers used in this work.

### LP benchmark

The first LP benchmark consisted of single-species FBA simulation with growth maximization. Seven genome-scale metabolic models (Table 2) were downloaded from the *BiGG Models*^14^ database. These models were selected to represent a wide variety of model sizes (uniformly distributed on a logarithmic scale). The simulations were repeated 10 times for each combination of model and solver to better estimate the computational time.

**Table 2:**
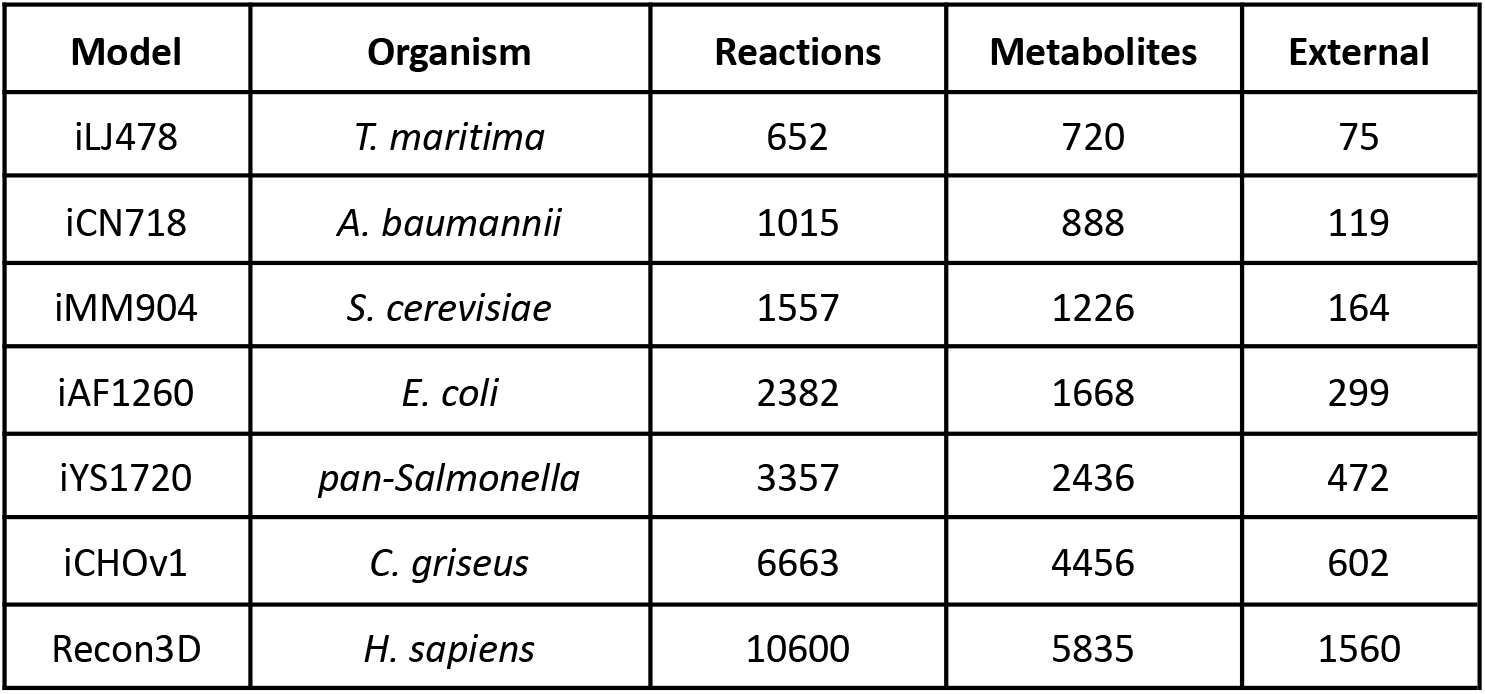
List of genome-scale metabolic models used in this work.

A second benchmark was performed for simulation of microbial communities with up to 20 members. The models were downloaded from the EMBL GEMs collection^6^ of 5587 bacterial models. For each community size, 10 community models were created by random sampling of species from the model collection. The communities were assembled using a compartmentalized approach and simulated by maximizing the sum of growth rates of all members.

### MILP benchmark

The MILP benchmark consisted on finding a minimal growth medium for each model in Table 2. The problem is implemented by minimizing the total number of active uptake reactions in each model subject to a non-zero growth rate. The number of binary variables corresponds to the number of external metabolites in each model. The computation was repeated 10 times for every combination of model and solver.

An additional robustness test was performed for solving MILPs with the Recon3D model using only a random sample of 312 external metabolites (i.e., 20% of the original number of binary variables). The test was performed 1000 times for each solver. In this case, due to their long computation times, GLPK and COIN were executed in a computing cluster with a configuration similar to the local environment (8 CPU cores with 16 GB of RAM).

## Results

### LP benchmark

Benchmarking results for solving LPs are presented in Figure 1. For single-species FBA, it can be observed that GUROBI was the fastest solver, followed by CPLEX, whereas SCIP was the slowest solver for models of small size. COIN starts with a performance comparable to HiGHS and GLPK, but it deteriorates for larger problem sizes, indicating some difficulty with scaling up. All problems are solved in a time-scale of milliseconds, with a negligible variation in runtime between replicates. The largest model (Recon3D), with approximately 10K variables and 6K constraints, can be solved in a range of 0.1 to 1.0 seconds.

**Figure 1:**
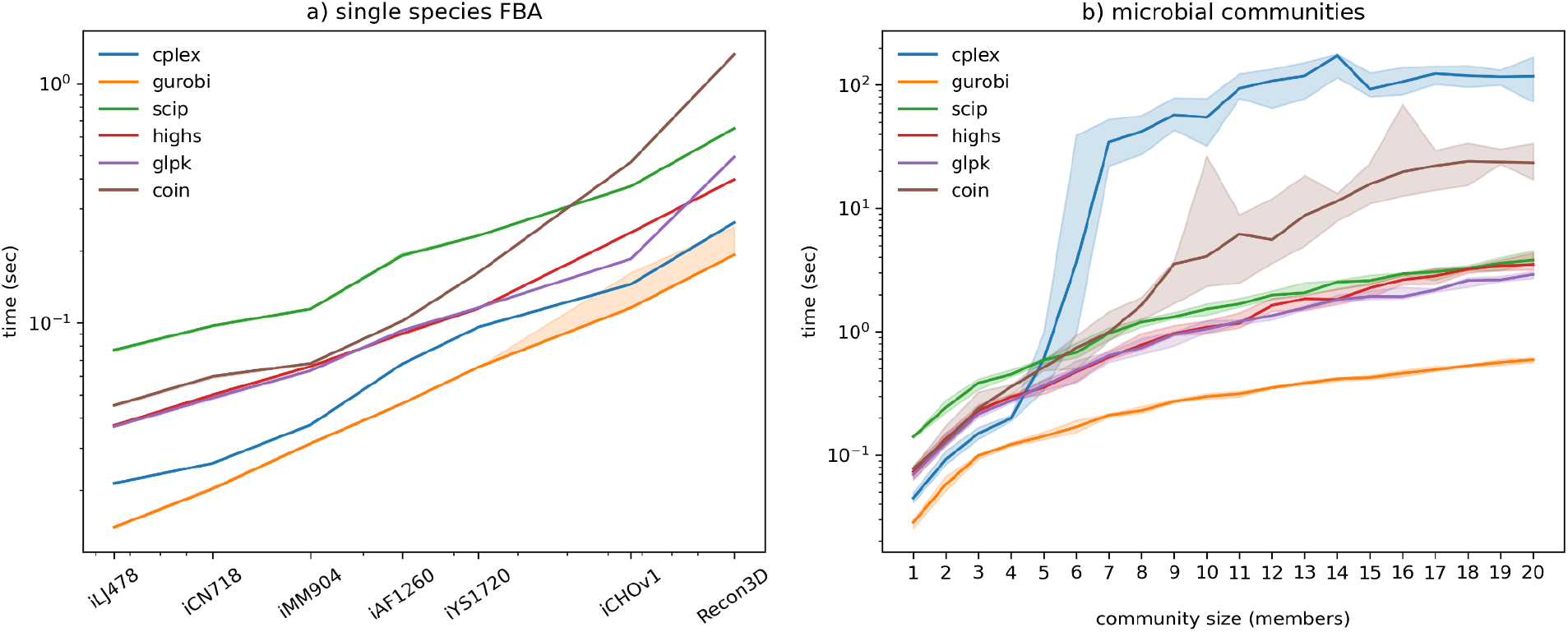
Benchmarking results for LP formulations. a) Single-species FBA. The horizontal axis (log-scale) represents the number of continuous variables (number of reactions in each model). b) Simulation of microbial communities. In both panels, lines represent the median of 10 replicates and the error bands represent the interquartile range.

For simulation of small microbial communities (up to 4-5 members), the results are comparable to single-species FBA. However, for larger community sizes, CPLEX suddenly decreases in performance. This change, by approximately 100-fold, is abrupt but stable, and could indicate some internal shift in heuristic strategies. GUROBI maintains its remarkable efficiency, with the exception of one outlier case. Three open-source solvers (SCIP, HIGHs, GLPK) perform very similarly, with a stable computational time that seems to approximate a complexity of *O(n*^*\**^*log(n))*.

### MILP benchmark

As expected, MILP problems have considerably higher runtimes than LP, in the order of seconds to minutes (Fig. 2a). The variation between runtimes (in the order of milliseconds) again becomes negligible. Again, GUROBI and CPLEX are the fastest solvers. In this case COIN is clearly the slowest solver, with a runtime, on average, one order of magnitude above the others. For smaller problems, GLPK has a runtime comparable to SCIP and HiGHS, but its performance deteriorates for larger problems, indicating issues with scalability. Both GLPK and COIN failed to solve the largest problem (Recon3D). Their execution was killed after one week of runtime. Interestingly, the runtime for solving MILPs did not increase monotonously with the size of the models. For instance, the iAF1260 model took longer to solve than the larger iYS17120 model, regardless of the choice of solver. This indicates that other factors (such as specific constraints resulting from network topology and stoichiometric coefficients) can contribute to the time required to find the optimal solution.

**Figure 2:**
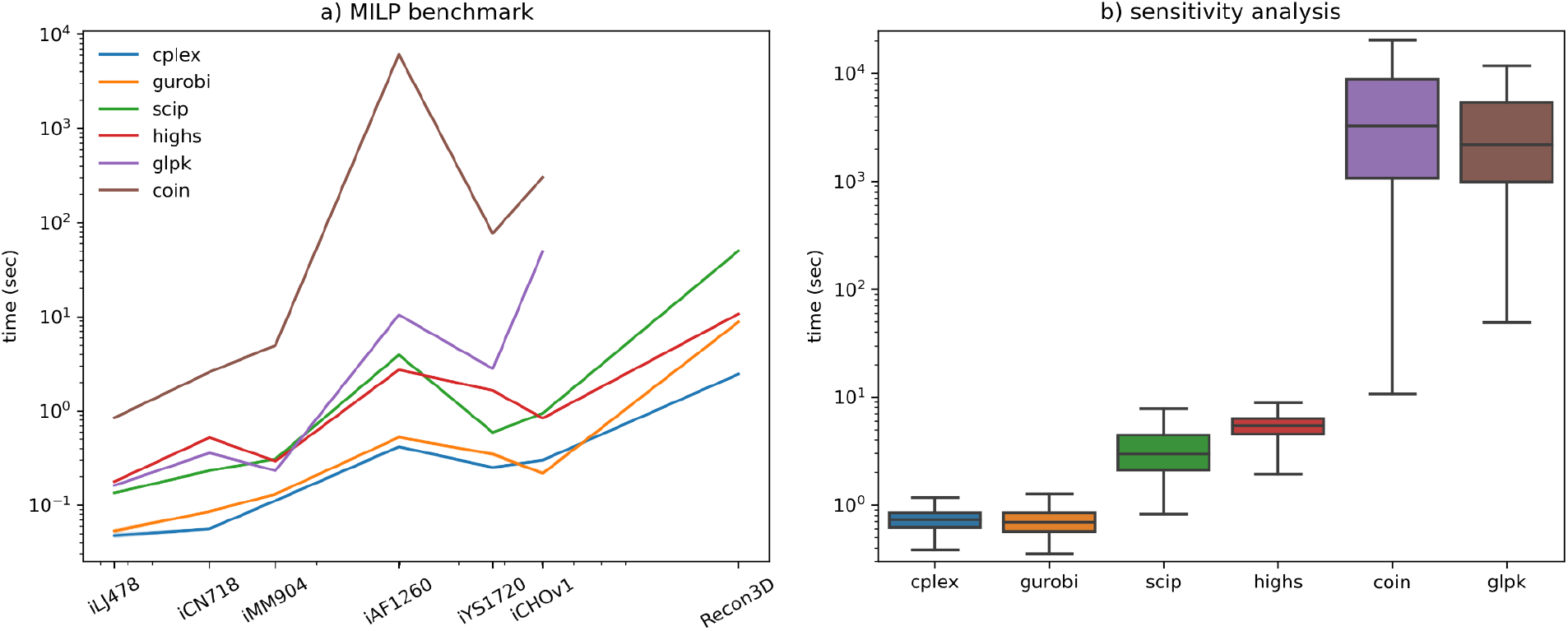
Benchmarking results for MILP formulations. a) Computation time for each model. The horizontal axis (log-scale) represents the number of binary variables (number of external metabolites in each model). Interquartile variation is represented by error bands but are too small to be detected. GLPK and COIN failed to solve the Recon3D model. b) Sensitivity analysis for Recon3D with random subsets of variables (1000 replicates).

To further analyze the robustness of the solvers towards perturbations in the MILP formulations, the Recon3D model was solved 1000 times with random subsets of 312 binary variables (i.e., considering only 20% of all uptake reactions). The reduced problem complexity allowed to obtain performance results for GLPK and COIN (Figure 2b). It can be observed that the solvers fall into three main categories: the fast solvers, CPLEX (0.74 ± 0.17 sec) and GUROBI (0.74 ± 0.24 sec); the intermediate, SCIP (3.53 ± 2.19 sec) and HiGHS (5.38 ± 1.48 sec); and the very slow GLPK (1.3 ± 2.1 h) and COIN (2.4 ± 4.0 h). It is quite clear that, despite the same problem size, the execution time can vary significantly, especially for the slowest solvers. This is likely influenced by the heuristics implemented by each solver for the *branch-and-cut* exploration of the MILP solution space.

## Discussion

Overall, this work shows that, although CPLEX and GUROBI are still the fastest solvers, some of the open-source solvers offer comparable performance. Even for the Recon3D model, with over 10K reactions, it is possible to run an FBA simulation in less than a second, and (with the exception of GLPK and COIN) it is possible to solve an MILP with over 1.5K binary variables in just a few seconds. Among the open-source solvers, GLPK and HiGHS performed the best for LP formulations, whereas SCIP and HiGHS performed better for MILP formulations. Hence, HiGHS might offer the best compromise between both types of problems. The sudden lack of performance of CPLEX for microbial community modeling was a surprising result.

In this work, I opted to run all solvers with their default parameter configurations. Each solver can be potentially fine-tuned by adjusting internal parameters (tolerances, pre-solving, heuristic search strategies, etc). Since each solver implements a different set of parameters, such fine-tuning is outside the scope of this work. Nonetheless, it was confirmed that all solvers returned the same objective values for every problem. The solution degeneracy typical of genome-scale models can, of course, result in alternative flux distributions with the same objective value.

Genome-scale metabolic modeling is a powerful framework to map genotypes to phenotypes and help to develop biotechnological solutions to address many of today’s societal challenges. As we try to understand the impact of environmental changes in the planetary microbiome or the impact of our own microbes and diets in our health, more detailed models and sophisticated simulation tools will emerge. The open-source solvers used in this work are currently available for genome-scale modeling, through a core simulation library^18^, and can be easily interfaced with in any other tool. This is a fundamental step for the ongoing integration of genome-scale modeling into complex bioinformatic workflows^19^ that will help untangle the metabolic complexity of our biosphere.

## Acknowledgements

This work was supported by ELIXIR Norway, funded by the Research Council of Norway, and the Cell4Chem project, funded under the 3rd ERA CoBioTech international joint call.

## Data availability

Source code and data are openly available at: https://github.com/cdanielmachado/solver_benchmark.

